# EEG and fMRI Evidence for Autobiographical Memory Reactivation in Empathy

**DOI:** 10.1101/715276

**Authors:** Federica Meconi, Juan Linde-Domingo, Catarina S. Ferreira, Sebastian Michelmann, Bernhard Staresina, Ian Apperly, Simon Hanslmayr

**Affiliations:** School of Psychology, University of Birmingham, B15 2TT.; Max Plank Institute Berlin for Human Development, 14195, Berlin; Princeton Neuroscience Institute, Princeton, NJ 08544.; Center for Human Brain Health, University of Birmingham, B15 2TT; Institute for Neuroscience and Psychology, University of Glasgow, G12 8QB

**Author notes:** **Corresponding author:** Federica Meconi (F.M.), School of Psychology, Hills Building, University of Birmingham, B15 2TT. Juan Linde-Domingo (J.L.D.), Max Plank Institute Berlin for Human Development, 14195, Berlin. Catarina S. Ferreira (C.S.F.), School of Psychology, Hills Building, University of Birmingham, B15 2TT. Sebastian Michelmann (S.M.), Princeton Neuroscience Institute, Princeton, NJ 08544. Bernhard Staresina (B.S.), School of Psychology and Center for Human Brain Health, Hills Building, University of Birmingham, B15 2TT. Ian A. Apperly (I.A.A.), School of Psychology, 52 Pritchatts Road, University of Birmingham, B15 2TT. Simon Hanslmayr (S.H.), Institute for Neuroscience and Psychology, 62 Hillhead Street, University of Glasgow, G12 8QB.

**Keywords:** Empathy, Autobiographical memory, EEG, fMRI, EEG pattern classifier

## Abstract

Empathy relies on the ability to mirror and to explicitly infer others’ inner states. Theoretical accounts suggest that memories play a role in empathy but direct evidence of a reactivation of autobiographical memories (AM) in empathy is yet to be shown. We addressed this question in two experiments. In experiment 1, electrophysiological activity (EEG) was recorded from 28 participants who performed an empathy task in which targets for empathy were depicted in contexts for which participants either did or did not have an AM, followed by a task that explicitly required memory retrieval of the AM and non-AM contexts. The retrieval task was implemented to extract the neural fingerprints of AM and non-AM contexts, which were then used to probe data from the empathy task. An EEG pattern classifier was trained and tested across tasks and showed evidence for AM reactivation when participants were preparing their judgement in the empathy task. Participants self-reported higher empathy for people depicted in situations they had experienced themselves as compared to situations they had not experienced. A second independent fMRI experiment replicated this behavioural finding and showed the predicted activation in the brain networks underlying both AM retrieval and empathy: precuneus, posterior parietal cortex, superior and inferior parietal lobule and superior frontal gyrus. Together, our study reports behavioural, electrophysiological and fMRI evidence that robustly supports the involvement of AM reactivation in empathy.

## 1. Introduction

When we encounter somebody who has a physical injury, like a broken leg, we feel we have a good understanding of their pain, especially if we have experienced that same injury in our life. Therefore, it is intuitively compelling to assume that empathy, i.e. the ability to share and understand others’ inner states, draws on first-hand experiences we collected in our own past, i.e. on autobiographical memories (AM). However, this compelling intuition cannot be taken for granted because empathy might instead be supported by semantic memory about the general experience of pain and the conditions in which it is likely to occur (Perry et al., 2011; Rabin and Rosenbaum, 2012). The present study used advanced imaging methods to distinguish carefully between the roles of autobiographical and semantic memory, and seek the first direct evidence of reactivation of AM in the service of empathy.

The claim for an interplay between AM and empathy is supported by several sources of convergent evidence. Healthy students show higher empathy levels for adults experiencing chronic pain if they can rely on their own general AM of physical pain when compared to control conditions (Bluck et al., 2013). Patients with congenital insensitivity to pain show attenuated self-rated empathy for others’ pain, for which they could have collected no experience or memory (Danziger et al., 2009). Several studies, provide evidence of common brain networks for AM retrieval and cognitive empathy (i.e. reasoning explicitly about mental states (Buckner and Carroll, 2007; Spreng et al., 2008; Spreng and Grady, 2009), with these brain areas including precuneus, posterior cingulate cortex (PCC), retrosplenial cortex, medial temporal lobe (MTL), temporoparietal junction (TPJ) and medial prefrontal cortex (mPFC, BA 10). Neuropsychological studies on patients with different kinds of memory impairments have reported generally convergent results about impoverished empathic abilities as measured by neuropsychological assessments or self-report questionnaires. Empathy failures are symptoms of several psychiatric disorders with long-term memory impairment, e.g. schizophrenia (Corcoran and Frith, 2003; Meconi et al., 2016). They are observed in patients with Alzheimer’s disease (Moreau et al., 2013; Ramanan et al., 2017), Korsakoff’s syndrome (Drost et al., 2019; Oosterman et al., 2011), Mild Cognitive Impairment (Moreau et al., 2015), Parkinson disease (Monetta et al., 2009; Pell et al., 2014; Xi et al., 2015), and Semantic dementia (Duval et al., 2012). In healthy ageing the affective side of empathy seems to decrease with age (Chen et al., 2014; Duval et al., 2011; Ze et al., 2014). However, conclusions from the latter studies must be treated with some caution because patients with a diagnosis involving different kinds of memory loss might show reduced empathy as a consequence of a global cognitive decline that impairs multiple executive functions that in turn show reduced self-reported empathy.

Curiously, the handful of studies on patients with amnesia, i.e. a memory disorder observed after focal hippocampal cortices damage that impairs the ability to consciously access AM, showed that empathy seems to be spared (Rosenbaum et al., 2007) or only mildly impaired (Beadle et al., 2013; Staniloiu et al., 2013), at least for cognitive empathy (“cold reasoning” about mental states) rather than affective empathy (“hot simulation” of other people’s states and experiences, Sessa et al., 2014b; Shamay-Tsoory et al., 2009). While studies of amnesia are potentially powerful sources of evidence evaluating the role of memory in empathy, they do not provide definitive evidence about the role of AMs. This is because the retrieval of AMs does not rely only on the hippocampal cortices but is underpinned by a network of brain areas that involves the prefrontal cortex and parietal areas including precuneus, posterior parietal cortex and the retrosplenial cortex (Boccia et al., 2019; Cabeza and St Jacques, 2007; Cotelli et al., 2012). In line with the systems consolidation account, by which memories become gradually independent of the hippocampus and stably stored in the neocortex, at least remote memories might still be available as a source of semantic knowledge or implicit memories for the patients (Antony et al., 2017; McClelland et al., 1995). This clearly leaves open the possibility that AMs are retrieved during empathy, and that this could be detected with appropriate methods.

Even though the studies mentioned above support the idea that empathy draws on AM, critical evidence is missing. In particular, it has not yet been demonstrated that AMs are actively retrieved in the service of empathy. In order to test for evidence of a re-activation of AM when empathizing with others, we investigated healthy adults’ empathy for an AM experience that the participants shared with the targets of empathy, in contrast with an un-shared experience for which participants had no AM. In light of evidence that the role of AM may be different for “affective” and “cognitive” empathy, we also varied whether the shared and un-shared experiences were emotionally potent (physical pain) versus emotionally neutral (e.g., visiting a museum).

Recent advances in multivariate pattern analysis methods show that brain activity patterns can be tracked during the encoding of new neutral episodes and re-observed during their retrieval (Linde-Domingo et al., 2019; Michelmann et al., 2016). Furthermore, recent neuroimaging studies show that reinstatement of autobiographical pain involves partial reinstatement of activity in the brain areas that process nociception (Fairhurst, Fairhurst, Berna, & Tracey, 2012; Forkmann, Wiech, Sommer, & Bingel, 2015). Therefore, we here tested for a direct evidence of online reactivation of AM when participants were required to explicitly rate their empathy awareness for others’ neutral and painful expriences in two independent experiments (Fig. 1c, d). In experiment 1, EEG was recorded during two sequential tasks: the first was a pain decision task, classically used to prompt an empathic reaction, in which targets for empathy were depicted in contexts for which participants either did or did not have an AM (Fig.1a); the second was a a task that explicitly required retrieval of the AM and non-AM contexts. The retrieval task was used to extract the neural fingerprints of AMs and non-AMs (Fig.1b). A linear discriminant analysis (LDA) EEG pattern classifier was trained during the retrieval task and tested on data obtained from the preceding empathy task to test for the online reactivation of the memories in explicit empathy. In experiment 2, fMRI was measured from an independent sample performing the same empathy task to test if it activated brain areas commonly associated with empathy and AM.

**Fig. 1.**
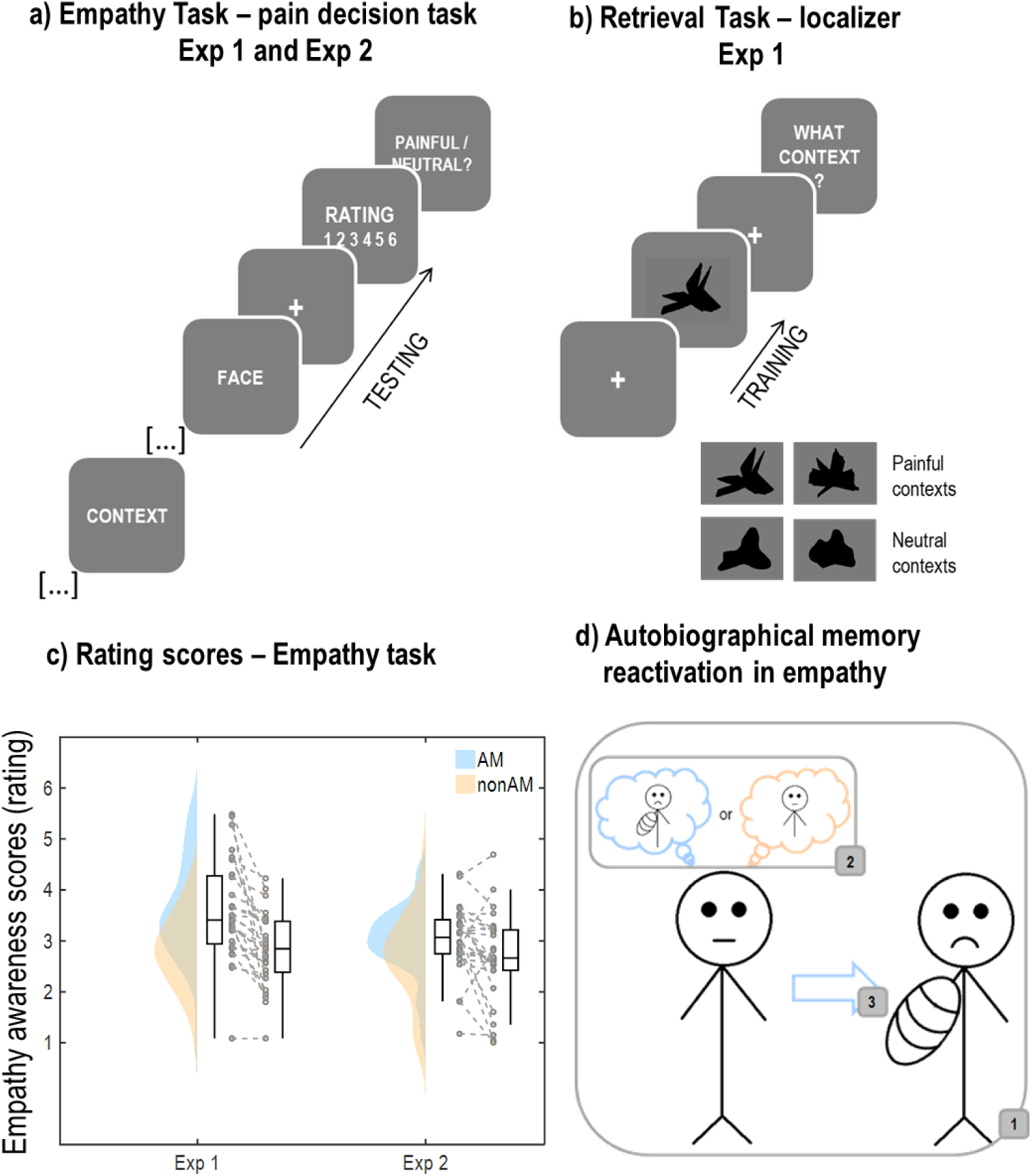
Experimental Design. a) Schematic representation of the empathy task used in experiment 1 and experiment 2. Participants were required to rate how much empathy they felt for the person depicted in the preceding context. b) Schematic representation of the retrieval task used in experiment1 that was used to train the LDA classifier. Participants first learnt to associate four abstract figures with the same sentences describing painful contexts presented during the empathy task (not shown here). In the actual task, for each trial participants were presented with one of the four figures and had to picture in their mind’s eye the context that they learnt to associate with that specific figure. c) Raincloud plots of the subjective reports of participants’ empathy awareness in experiment 1 and experiment 2. d) Concept of the study; when we encounter someone who shares our same physically painful experience, memory of that experience is reactivated to empathize.

**Fig. 2.**
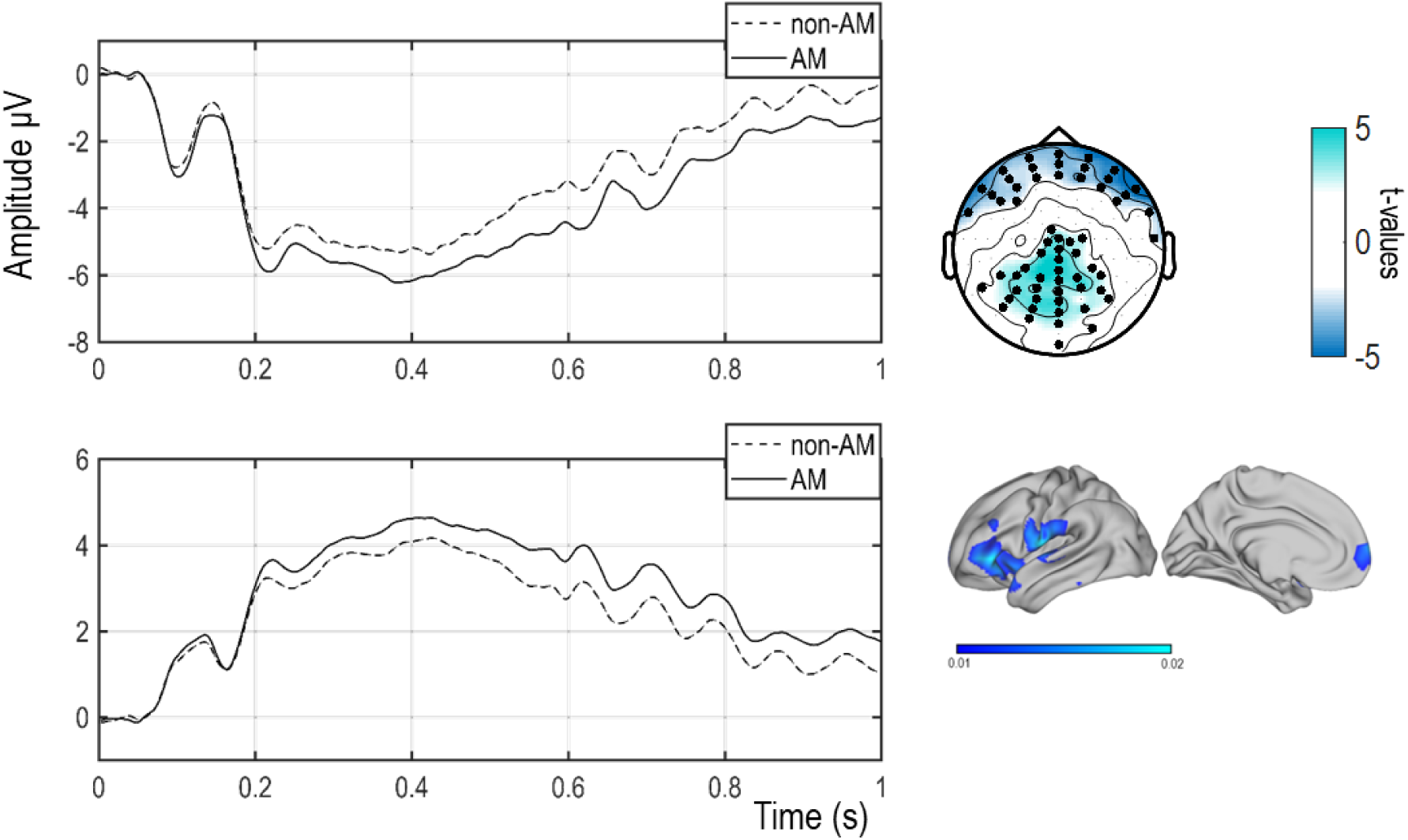
ERPs results. Left panel: ERPs time-locked to the onset of the face and reflecting AM and non-AM at the anterior (upper panel) and the posterior cluster (bottom panel. Right top panel: clusters analysis performed over all the electrodes in a 0 – 1 sec time-window. Colors code t-values. Right bottom panel: source localization of the AM vs non-AM contrast.

**Fig.3.**
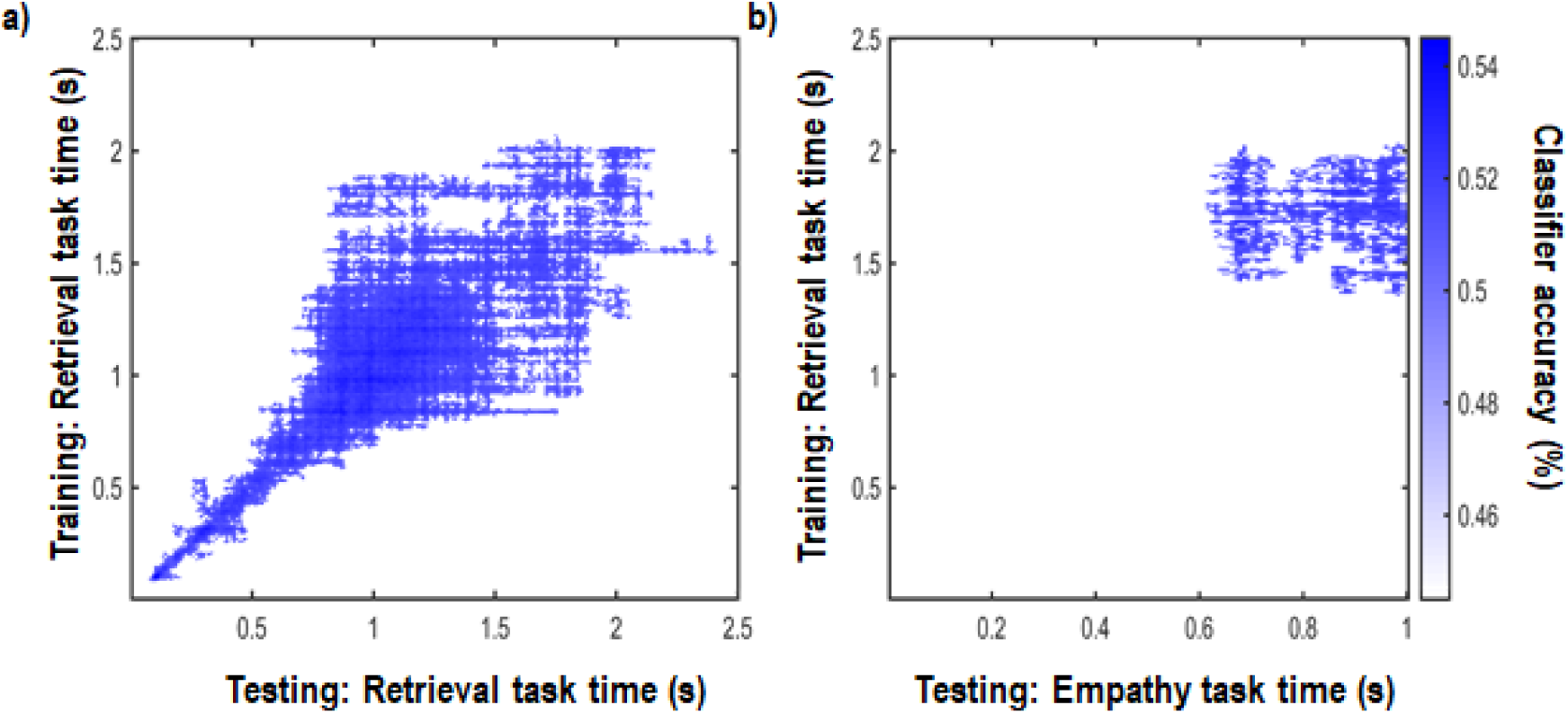
LDA results. a) Sanity check: time by time generalization matrix showing significant classification of AM vs non-Am within the retrieval task. b) Time by time generalization matrix (i.e. training and testing at each time-point) showing significant classification of AM vs non-AM across tasks.

**Fig. 4.**
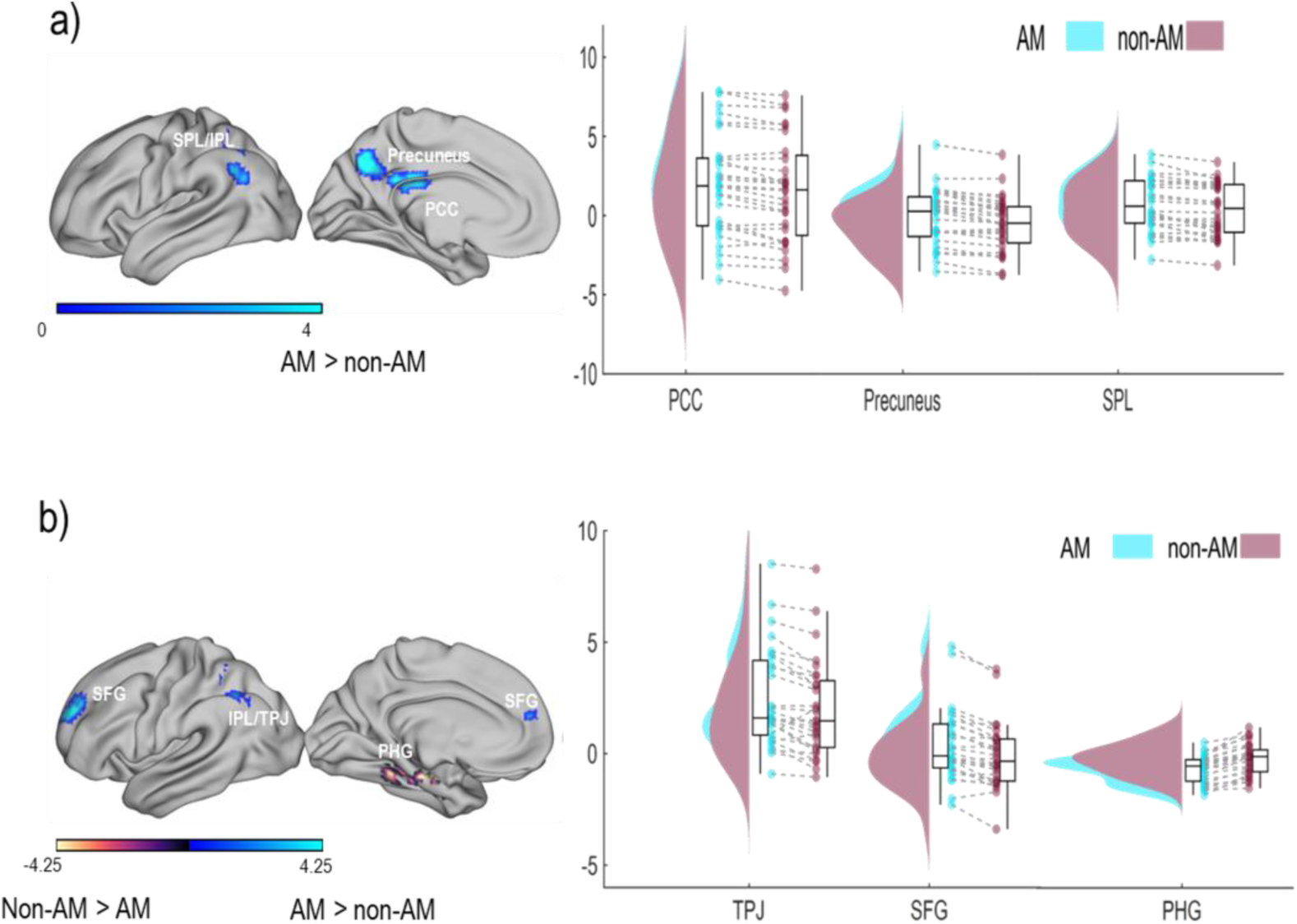
FMRI results. Whole-brain analysis results (left panel) and raincloud plots (right panel) of the activation in each condition and each cluster. a) Whole-brain analysis related to the presentation of the context. Only the contrast AM>non-AM showed significant clusters. b) Whole-brain analysis related to the presentation of the face. Figure shows significant clusters resulting from both the AM>non-AM and non-AM>AM contrasts.

## 2. Material and Methods

The protocol for both experiments was approved by the University of Birmingham Research Ethics Committee (ERN-16-0101A). Written informed consent was obtained from all participants for both experiments.

### 2.1. Participants

We aimed for a sample size of 28 participants. This is consistent with previous studies using the pattern classifier in the field of memory (Linde-Domingo et al., 2019; Michelmann et al., 2016). Also, using the PANGEA analysis tool (https://jakewestfall.shinyapps.io/pangea/), 28 participants was judged to be able to detect the main effect and the 3-way interaction of a medium effect size with a power exceeding 0.9. All the participants, for both experiments, were recruited through the Research Participant Scheme of University of Birmingham for cash (£10/h) or course credits (1 credit/h). All of the participants had normal or corrected-to normal vision. The eligibility criteria included native or excellent English proficiency, no history of neurological or psychiatric disorder, and having an experience of intense physical pain in their past. In order to ascertain that all the eligibility criteria were met, students who signed up for the study were contacted and screened before they were accepted as participants for the studies. During this initial screening phase, students were asked about their English proficiency, they had to complete a questionnaire were pathological history or psychotropic drugs assumptions were checked and they had to complete the Autobiographical Memory Questionnaire, AMQ (Rubin et al., 2003), for an experience of intense physical pain and for one emotionally and physically neutral that they were asked to report. Students who could not report any experience of intense physical pain or of a neutral experience could not be participants in these studies.

#### 2.1.1. Experiment 1. EEG

Thirty-five healthy students took part in the experiment (mean age = 22, SD = 5). Four were left-handed, 6 were males. Seven were discarded from the final sample, three were not Caucasian (we showed pictures of Caucasian faces and previous studies have shown that empathic responses are subject to ethnicity bias (e.g., Sheng & Han, 2012), two could not complete the task due to equipment failure and two for too low number of trials due to inaccurate responses. The final sample was composed of 28 participants (mean age = 21.96, SD = 4.82), four were males and four left-handed.

#### 2.1.2. Experiment 2. fMRI

Thirty three healthy students took part in the experiment (mean age = 25, SD = 5.9). Participants were all right-handed, 15 were males. Five were discarded from the final sample, two served as pilots to adjust the timing of the paradigm and make it suitable for the fMRI environment; one participant could not complete the acquisition session in the scanner, two were discarded for excessive movements (more than 1 voxel size, 3mm). The final sample was composed of 28 participants (mean age = 24.71, SD = 5.86), eleven were males.

### 2.2. Questionnaires

As mentioned in section 2.1, before accepting students as participants for the study, the students who signed up for the study underwent a screening phase. The screening consisted of collecting information about students’ history of pathological morbididy, English proficiency and, critically, an AM of intense physical pain and of a neutral experience, in terms of emotion and pain. Candidates were therefore asked to report these AMs and complete the AMQ for both autobiographical episodes. The AMs reported by the participants were on average 4.65 years old (SD = 5 y) for experiment 1; 4.46 years old (SD = 5.72 y) for experiment 2.

At the end of the experimental session, dispositional empathy resources and the ability to recognize and describe participants’ own emotions were assessed with the Empathy Quotient, EQ, and the Interpersonal Reactivity Index, IRI (Baron-Cohen and Wheelwright, 2004; Davis, 1983) and with the Toronto Alexithymia scale, TAS-20 (Bagby et al., 1994), respectively. Participants from both experiments fell in the normal range of the EQ (experiment 1: M = 51.14 SD = 9.99; experiment 2: M = 46.89 SD = 12.37), and had on average normal ability to describe their emotions as showed by the TAS score (experiment 1: M = 45.96, SD = 12.22; Exp2: M = 43.96, SD = 12.01). The IRI scores for both experiments are reported in Table 1. These measures were also used to explore any relation with the neural responses but no correlation was found significant (for further details on the correlation analysis please see the Supplementary Materials).

**Table 1.** Questionnaires. Participants’ scores to the Interpersonal Reactivity Index in both experiments. T-tests are performed on the two independent samples.

|  | Exp 1 |  | Exp 2 |  | *t*(27) | *p* |
| --- | --- | --- | --- | --- | --- | --- |
|  | Mean | SD | Mean | SD |  |  |
| F | 3.89 | 0.89 | 3.81 | 0.71 | 0.569 | >0.5 |
| Pt | 3.74 | 0.75 | 3.73 | 0.81 | 0.067 | >0.5 |
| EC | 4.2 | 0.61 | 3.86 | 0.85 | 2.135 | 0.042 |
| Dp | 2.92 | 0.75 | 2.85 | 0.74 | 0.51 | >0.5 |

### 2.3. Stimuli and Procedure

All the stimuli were presented on a grey background of a 17’’ computer screen with a refreshing rate of 70 Hz. The tasks were programmed using Psychtoolbox.

#### 2.3.1. Experiment 1: EEG

Participants performed two tasks in the same experimental session. For all participants, the first of these tasks was the empathy task used in previous studies (Meconi et al., 2018; Sessa et al., 2014b) and the second one was a retrieval task (Fig.1a and b).

##### 2.3.1.1. The empathy task: stimuli and procedure

The stimuli for the empathy task were sentences, describing specific contexts, followed by faces, the task required to rate participants’ empathy for the person as depicted in the preceding context. The faces were a set of 16 identities, 8 males and 8 females with a painful or a neutral facial expression. The faces were in shades of grey and they were equalized for luminance with the SHINE toolbox (Willenbockel et al., 2010). The sentences described contexts of a person feeling physical pain or depicted in an emotionally neutral context. The critical manipulation in this task was that the targets of participants’ empathy were depicted in contexts for which they had or had not a related AM. Therefore, two contexts (one describing physical pain and the neutral one) were taken from participants’ autobiographical experience. In order to tailor the contexts for each participant, we screened them prior to the experimental session as soon as they signed up for the study. Participants were asked to report an experience of intense physical pain and an emotionally and physically neutral experience for which they completed the AMQ. A physically painful and a neutral context that didn’t belong to participants’ autobiographical experiences were also identified and used for the non-AM contexts. The four contexts identified for each participant were described in the empathy task by a sentence that always followed the structure “This person got – […]” or “This person did – […]” so that all the sentences had the same syntactic complexity.

Each trial started with a fixation cross (600 ms). Participants were then presented with the sequence of a sentence (3 s) and a face (500 ms) interleaved by a variable fixation cross (800 −1600 ms jittered in steps of 100 ms). The task was to subjectively rate on a scale of 6 points how much empathy participants felt for the person as depicted in the presented context. The rate was self-paced and presented after another fixation cross that was on the screen for 500 ms. At the very end of the trial, participants were asked to indicate whether the face, regardless of the context, had a painful or a neutral expression. The task was composed of 48 trials per condition that were pseudo-randomized in a way that the conditions were balanced over the full session. There were 192 trials in total subdivided in 4 blocks. An illustration of the task is shown in Fig.1a.

##### 2.3.1.2. The retrieval task: Stimuli and procedure

For the retrieval task, 16 shapes were created ad hoc in total. For each participant a unique subset of 4 of these shapes was presented. The first step was to generate 8 random polygons with equal number of black pixels. The polygons were then blurred with a Gaussian filter and all the pixels in shades of grey were made black to create 8 rounded shapes. As a last step, the number of black pixels was equalized across all the shapes (i.e., random polygons and the rounded shapes). The shapes were shown on a grey background (Fig.1b).

In the retrieval task, participants were required to picture in their minds’ eye the contexts described in the empathy task. This task acted for the EEG pattern classifier as a localizer to extract the neural fingerprints of the contexts to then probe the data from the empathy task. Therefore, to avoid perceptual confounds when applying the classifier across tasks, participants were presented with their unique subset of four shapes that were used to cue the contexts described in the empathy task. Before starting the retrieval task, participants underwent a “learning phase” in which they learnt to associate each shape with one of the contexts, each sentence-shape pair was presented twice before memory for the associations was tested. The retrieval task could only start when full accuracy was reached in the “learning phase”. To test memory for the sentence-shape association, participants were presented only with a figure at a time and had to indicate what was the context associated with the shape. One cue-word per sentence was chosen to cue to the related context and allow responses (e.g., we used “arm” as a cue-word for the sentence “This person got their right arm broken”; “ligament” for “This person got their ligament torn”; “Museum” for “This person visited the Birmingham Museum of Art” and “laptop” for “This person bought a new laptop” and so forth). The four cue-words were placed equally spaced horizontally at the center of the screen and their order was randomized with a Latin square in such a way that each word had the same likelihood to appear at one of the four locations (e.g., “arm” “ligament” “museum” “laptop”; “museum” “arm” “laptop” “ligament” etc.). Participants could press one of four keys on the computer keyboard that spatially corresponded to the location of the cue-word (“d” for the cue-word appearing on the very left, “f” for the cue-word appearing the central left, “j” for the one at the central right and “k” for the one at the very right location). The memory test could end after 8 correct answers, i.e. 2 times each pair. One error within a block of 8 trials would be followed by the repetition of a new block of 8 trials, until 100% accuracy would be reached. Once the memory associations test was successful participants could start the practice session of the retrieval task. Participants could familiarize with the retrieval task with a block of 8 trials that could be repeated until they felt confident they understood the task.

In the retrieval task, participants were only shown the shapes. In each trial, one shape was shown for 3 secs and participants had to picture in their mind’s eye the context associated with that shape. Within the time the shape was on the screen, they were required to rate the vividness of the context as soon as they could picture it in their mind’s eye, by pressing one of six response keys “s”, “d”, “f”, “j”, “k”, “l”, with “s” for “not vivid at all” to “l” for “very vivid”. If they did not press any button, a “No Response” was recorded and the trial excluded from the analysis. Participants were then asked to indicate which context they saw in their mind’s eye. They could answer in the same way as they did for the memory association test with the further option that if they could not remember what context was associated with that shape, they could press the space bar for “forgotten” and move to the next trial. Responses were not time-pressured but only correct trials were included in the analysis. There were 60 trials per condition that were pseudo-randomized to balance the distribution of all the conditions over the total of 240 trials that constituted the full session. The task, depicted in Fig.1b, right panel, was subdivided in 4 blocks.

#### 2.3.2. Experiment 2: fMRI

The screening phase, the questionnaires and the procedure were the same as those used in experiment 1 with the exception of the necessary adjustments in the timing of the events applied to the empathy task in order to make it suitable for the fMRI environment (for additional details, see Supplementary Material S1.3.1.).

### 2.4. Data acquisition and analysis

#### 2.4.1. Experiment 1: EEG

The EEG was recorded using a BioSemi Active-Two system from 128 Ag/AgCl active electrodes. The EEG was re-referenced offline to the average reference. Three additional external electrodes were placed below the left eye and on the lateral canthi of each eye to record vertical electroculogram (EOG). EEG and vertical EOG signals were digitized at a sampling rate of 1024 Hz via ActiView recording software (BioSemi, Amsterdam, the Netherlands).

EEG data was analyzed with MATLAB (©Mathworks, Munich, Germany) using the open-source FieldTrip toolbox (http://fieldtrip.fcdonders.nl/) and in-house Matlab routines.

##### 2.4.1.1. Preprocessing

###### 2.4.1.1.1. The Empathy task

EEG data was first segmented into epochs of 2 s, starting 1 s before the onset of the face. The epoched data was visually inspected to discard large artifacts from further analysis. Further preprocessing steps included Independent Component Analysis for ocular artifacts correction and re-referencing to average reference. After removing trials which were contaminated by eye and muscle artefacts, an average of 45 trials (range: 34-48) remained for AM and 45 trials (range: 37-48) for non-AM condition.

###### 2.4.1.1.2. The Retrieval task

EEG data were first segmented into epochs of 4 s, starting 1 s before the onset of the cue. The epoched data was visually inspected to discard large artifacts from further analysis. Further preprocessing steps included Independent Component Analysis for ocular artifacts correction and re-referencing to average reference. After removing trials which were contaminated by eye and muscle artefacts, an average of 51 trials (range: 42-60) remained for AM and 50 trials (range: 38-58) for non-AM condition.

##### 2.4.1.2. Event-related potentials (ERPs) analysis: the Empathy task

ERPs were time-locked to the onset of the face. We computed ERPs in response to painful and neutral faces. To test any involvement of memory in the empathy task, we contrasted ERPs time-locked to the onset of the faces reflecting the processing of the preceding context (AM vs. non-AM). To check whether there was any difference between emotional content of the memories we also contrasted painful and neutral memories separately for AM and non-AM.

##### 2.4.1.3. Linear Discriminant Analysis EEG pattern classifier

Linear Discriminant Analysis (LDA) is a multivariate pattern analysis method that finds a decision boundary that allows distinguishing the pattern of brain activity associated with one category of stimuli from the pattern of brain activity that is associated with another category of stimuli. This is based on specified features of the EEG signal. It can then estimate with certain accuracy whether the pattern of brain activity in data that was not used to find the decision boundary, is more similar to one or the other category of stimuli.

In order to reduce unwanted noise and computational time, the signal was filtered between 0.1 and 40 Hz and down sampled to 128Hz before classification with a baseline correction window of 500ms before the onset of the stimuli.

The LDA was trained and tested on the EEG raw patterns (i.e. amplitude of the signal on each of the 128 electrodes), for each participant and at each time point and regularized with shrinkage (Blankertz et al., 2011). To make sure that the output was not biased by the signal to noise ratio due to the different amount of trials, we equalized the number of trials for AM and non-AM before train the classifier.

The classifier was trained on the raw signal (i.e. amplitude of EEG on each electrode) acquired while participants were performing the retrieval task in the time-window including the presentation of the cue to detect systematic differences between the EEG patterns reflecting the representation of AM and non-AM contexts. It was then tested on the signal independently acquired while participants were performing the preceding empathy task in the time-window from the onset of the face until the rating was made. The aim of the LDA was to test for the online reactivation of the memory in preparation of the explicit judgement of participants’ empathy awareness.

Before training and testing the LDA in two different datasets, we trained and tested the classifier on the retrieval task during the presentation of the cue to show that the task was successful to act as a localizer of the representation of the AM and non-AM contexts. A K-fold cross-validation procedure with 5 repetitions was used to train and test the classifier. The output of this analysis is the accuracy with which the classifier could distinguish between the two memory contexts for each time-point over all trials and electrodes. Therefore, the LDA reduces the data into a single decoding time course per dataset.

##### 2.4.1.4. Source Analysis

A standardized boundary element model was used for source modelling, which was derived from an averaged T1-weighted MRI dataset (MNI, www.mni.mcgill.ca). That was used in combination with individual electrode positions. Individual electrodes’ coordinates were logged with a Polhemus FASTRAK device (Colchester, Vermont, USA) in combination with Brainstorm implemented in MATLAB 2014b (MathWorks). For three participants the standard electrode coordinates were used due to technical problems during the experimental session.

For source reconstruction, a time-domain adaptive spatial filtering linear constrained minimum variance (LCMV) beamformer (Van Veen et al., 1997), as implemented in fieldtrip was applied. Source analysis was carried out for the time-domain ERP components that revealed significant results on the scalp level.

##### 2.4.1.5. Statistical analysis

###### 2.4.1.5.1. Behaviour: the Empathy Task

Mean proportions of accurate responses given within +/-2.5 SD from the average reaction time of each participant and mean proportions of the empathy awareness scores were computed for each condition and inserted in two repeated measures ANOVAs with a 2 (Emotional memory: Painful vs. Neutral) x 2 (Memory: Autobiographical vs. non-Autobiographical) x 2 (Facial expression: Painful vs. Neutral) as within-subject factors. Bonferroni corrected paired-sample t-tests were conducted when appropriate to explore significant interactive effects. Partial eta squared (*η*_*p*_^*2*^) are reported for completeness and transparency. Effect sizes are reported as eta squared (*η*^*2*^) calculated as the ratio between the sum of squares of each effect and the sum of the sum of squares of all the effects and their errors, 95% confidence intervals (CI) of the mean differences between conditions are reported in squared brackets.

###### 2.4.1.5.2. ERPs

Cluster-based permutation tests were performed over the whole scalp and over a 1sec time-window on the event-related potentials time-locked to the onset of the face. We tested for significant differences between painful and neutral facial expressions in order to replicate previous findings and show an ERP empathic response to faces. Additionally, preliminary analysis was carried out to test for any involvement of memory in the pain decision task and whether there was any difference related to the emotional content of the memory. To this end, cluster-based permutation tests were performed on the ERPs time-locked to the onset of the face regardless of the facial expression contrasting AM vs. non-AM and painful and neutral contexts separately for AM and non-AM.

###### 2.4.1.5.3. LDA classifier

For the classifier analysis, an empirical null distribution was created with a combined permutation and bootstrapping approach (Stelzer et al., 2013) that tested whether the maximum cluster of accuracy values above the chance level was statistically significant. Clusters were identified on the basis of the number of adjacent pixels found with the matlab function bwlabel. We used the LDA in 100 matrices with pseudo-randomly shuffled labels independently for each participant and created a null distribution of accuracy values that we contrasted with the LDA outputs obtained with the real data. This was done by sampling with replacement 100000 times from the real and random data of each subject and computing a group average. This procedure resulted in an empirical chance distribution, which allowed us to investigate whether the results from the real-labels classification had a low probability of being obtained due to chance (p < 0.05) (i.e., exceeding the 95^th^ percentile).

#### 2.4.2. Experiment 2. fMRI

Data acquisition was performed with 3T Philips Medical Systems Achiva MRI scanner using a 32-channel head coil. Functional T2-weighted images were acquired with isotropic voxels of 3mm, repetition time [TR] = 1750 msec, echo time [TE] = 30 msec, field of view [FOV] = 240×240×123 mm, and flip angle = 78°. Each volume comprised 33 sequentially ascending axial slices with an interslice gap of 0.75mm). Each participant underwent four blocks of scan series, one full block comprised 410 volumes. A high-resolution T1-weighted anatomical scan was acquired with an MPRAGE sequence (TR = 7.4 msec, TE = 3.5 msec, isotropic voxel size of 1mm, FOV = 256×256×176, flip angle = 7°) after the first two functional scanning blocks. The MR scanner was allowed to reach a steady state by discarding the first three volumes in each of the four scan series block.

##### 2.4.2.1. Preprocessing

The analyses were performed using the SPM12 toolbox (University College London, London, UK; http://www.fil.ion.ucl.ac.uk/spm/). For each scanning block, a motion realignment of each slice to the first slice was carried out before time realignment (slices corrected to the middle one). Data was then linearly detrended, using a Linear Model of Global Signal algorithm (Macey et al., 2004) to remove any minimal fluctuation due to the physical setting. Functional images served as reference for the co-registration with the anatomical image. The data was further normalized to an MNI template, and finally, images were spatially smoothed with an 8-mm FWHM Gaussian kernel.

##### 2.4.2.2. Whole-brain analysis

Two separate univariate analyses were carried out for two different time-windows, one analysis was time-locked to the onset of the face, and the other was time-locked to the onset of the context. This was only done to parallel experiment 1 and not to test for any functional dissociation between the two time-windows. In both cases statistical parametric maps were created for each participant’s block of trials.

AM and non-AM conditions were directly contrasted in paired-sample t-tests on a group-level analysis.

The first analysed fMRI data were time-locked to the onset of the context. Regressors were defined for AM and non-AM related to the onset of the contexts regardless of the emotional content of the context described by the sentence. Additional regressors of no interest were again included in the design matrix to explain variance in the data not due to the experimental manipulation under investigation and the 6 motion parameters obtained during the realignment phase of the preprocessing. Sixty statistical parametric maps were created (4 blocks × 15 regressors) for each participant.

The second analysed fMRI data were time-locked to the onset of the face. Regressors were defined for autobiographical and non-autobiographical memories time-locked to the onset of the faces regardless of their emotional expression. Additional regressors of no interest were included in the design matrix to explain variance in the data not due to the experimental manipulation under investigation plus the 6 motion parameters. Fifty-four statistical parametric maps were created (4 blocks × 14 regressors) for each participant. Additional information on the regressors of no interest are reported in Supplementary Material S1.3.2.2.

##### 2.4.2.3. Statistical analysis

###### 2.4.2.3.1. Behavior

Mean proportions of the empathy awareness scores were computed for each condition and inserted into a repeated measures ANOVA with a 2 (Emotional memory: Painful vs. Neutral) x 2 (Memory: Autobiographical vs. non-Autobiographical) x 2 (Facial expression: Painful vs. Neutral) as within-subject factors. Bonferroni corrected paired-sample t-tests were conducted when appropriate to explore significant interactive effects. Partial eta squared are reported for completeness and transparency. Effect sizes are reported as eta squared (*η*^*2*^) calculated as the ratio between the sum of squares of each effect and the sum of the sum of squares of all the effects and their errors, 95% confidence intervals (CI) of the mean differences between conditions are reported in squared brackets.

###### 2.4.2.3.2. fMRI

For both time-windows, a within-subject analysis was carried out on the data set of each participant to obtain the mean statistical parametric map for each experimental condition. Finally, a group-level paired-sample t-test contrasting AM and non-AM was performed. A cluster-wise analysis was performed with uncorrected *p* = .001 and then Family Wise Error correction was applied for multiple comparison (cluster *p* threshold = .05). Peak voxel MNI are reported in brackets. Further information on the fMRI analysis and results can be found in the Supplementary Materials (S1.3.2.1 and S2.2.1).

## 3. Results

### 3.1. Behavioural results

#### 3.1.1. Experiment 1: EEG

Individual scores of the empathy awareness revealed a main effect of the type of memory *F*(1,27) = 22.319; *p* = .000064, *η*_*p*_^*2*^ = .453, *η*^*2*^ = .092, M_diff_ = .767 CI_95_ = [.434 1.10] such that AM context induced higher empathy rates than non-AM contexts; of the emotional content of the memory *F*(1,27) = 50.902; *p* < .000001, *η*_*p*_^*2*^ = .653, *η*^*2*^ = .157, M_diff_ = 1.0, CI_95_ = [−1.288 −.713] and of the facial expression *F*(1,27) = 42.270; *p* < .000001, *η*_*p*_^*2*^ = .613, *η*^*2*^ = .193, M_diff_ = 1.108, CI_95_ = [−1.456 −.760] such that painful conditions induced higher rates than neutral conditions. The two-way interaction between emotional content of the memory and of the facial expression was significant *F*(1,27) = 18.390; *p* = .000206, *η*_*p*_^*2*^ = .405, *η*^*2*^ = .08. Further exploration of the two-way interaction revealed that painful faces drove higher rates of empathy awareness than neutral faces when the emotional content of the preceding memory was painful (*t*(27) = 9.7, *p*_*c*_ < .0000001, M_diff_ = 1.821, CI_95_ = [1.436 2.207]) but not when it was neutral (*p*_*c*_ = .167). In the same vein, painful memories reported higher empathy awareness scores than neutral memories when followed by painful (*t*(27) = 10.825, *p*_*c*_ < .0000001, M_diff_ = 1.714, CI_95_ = [1.389 2.038]) but not neutral faces (*p*_*c*_ = .286). The three-way interaction between the three factors was also significant *F*(1,27) = 11.002; *p* = .003, *η*_*p*_^*2*^ = .290, *η*^*2*^ = .003 such that empathy rates for painful faces was higher for painful when compared to neutral contexts for both AM (*t*(27) = 8.219, *p*_*c*_ < .0001, M_diff_ = 1.927, CI_95_ = [1.165 2.689]) and non-AM (*t*(27) = 6.397, *p*_*c*_ < .0001, M_diff_ = 1.500, CI_95_ = [.738 2.262]) but this difference was higher for AM than non-AM contexts (*t*(27) = 2.122, *p* = .043, M_diff_ = .427, CI_95_ = [.014 .840]).

#### 3.1.2. Experiment 2: fMRI

Individual scores of the empathy awareness revealed a main effect of the type of memory *F*(1,27) = 7.210; *p* = .012, *η*_*p*_^*2*^ = .211, *η*^*2*^ = .021, M_diff_ = .355, CI_95_ = [.084 .626]; of the emotional content of the memory *F*(1,27) = 48.860; *p* < .000001, *η*_*p*_^*2*^ = .644, *η*^*2*^ = 0.186, M_diff_ = 1.064, CI_95_ = [.752 1.376] and of the facial expression *F*(1,27) = 38.863; *p* = .000001, *η*_*p*_^*2*^ = .590, *η*^*2*^ = 0.186, M_diff_ = 1.065, CI_95_ = [.714 1.415]. The two-way interaction between emotional content of the memory and of the facial expression was significant *F*(1,27) = 19.995; *p* = .000126, *η*_*p*_^*2*^ = .405, *η*^*2*^ = .096, so was the one between the emotional content and the type of memory *F*(1,27) = 4.758; *p* = .038, *η*_*p*_^*2*^ = .150, *η*^*2*^ = .008. Further exploration of the two-way interactions revealed that painful faces drove higher rates of empathy awareness than neutral faces when the emotional content of the preceding memory was painful (*t*(27) = 8.8, *p*_*c*_ < .0000001, M_diff_ = 1.83, CI_95_ = [1.404 2.257]) but not when it was neutral (*p*_*c*_ = .280). In the same vein, AM reported higher empathy awareness scores than non-AM when they were painful (*t*(27) = 3.016, *p*_*c*_ = .006, M_diff_ = .578, CI_95_ = [.185 .971]) but not when they were neutral (*p*_*c*_ = .348). However, painful memories reported higher scores of empathy than neutral memories, were they either autobiographical or not (min *t*(27) = 5.17, *p*_*c*_ = .00002, M_diff_ = .841, CI_95_ = [.507 .1.175]). The three-way interaction did not reach significance level *F*(1,27) = 2.294; *p* = .142, *η*_*p*_^*2*^ = .078, *η*^*2*^ = .0006.

The behavioural results from the two experiments showed that individuals depicted in contexts describing participants’ autobiographical contexts drove enhanced explicit judgements of empathy awareness when compared to contexts describing non-autobiographical contexts independently of all the other factors. These results are shown in Fig.1c).

### 3.2. EEG results

#### 3.2.1. ERPs and source analysis

Cluster analysis conducted over a 1sec time-window, from the onset of the face until the presentation of the rating, revealed one anterior and one posterior cluster of electrodes showing that ERPs significantly differ as a function of the type of memory (anterior: p = 0.002; posterior: p = 0.002). Fig.2 depicts ERPs for AM and non-AM in the left panel and the topography of the significant clusters in right upper panel, t-values are plotted). Source analysis estimated that the neural source of this effect was the Superior Frontal Gyrus, BA 10, MNI: [−10 69 0] (Fig.2 right bottom panel). No difference was found for separate contrasts between emotional contents of the context for neither AM (p = .06) nor for non-AM (p = .18), therefore we did not further analyze differences between emotional contents of the contexts.

Additionally, in line with previous studies on empathy for physical pain, cluster analysis also revealed a classic ERP response associated with empathic processes (e.g., Sessa, Meconi, & Han, 2014), i.e. painful faces elicited more positive ERP responses than neutral faces (p = 0.004). Consistently, source analysis estimated that the neural source of this effect was the Inferior Frontal Gyrus, BA 9, MNI: [−62 21 30] and the Parietal Lobule, BA 7, MNI [30 −69 48].

#### 3.2.2. LDA

We first ran a sanity check of the classifier on the retrieval task. The classifier was trained and tested with a K-fold cross-validation procedure during the presentation of the cue (Fig.1b) as reported in section 2.4.1.3. Since we were interested in investigating whether AM is reactivated in empathy, we checked that the classifier could distinguish first of all whether the context pictured in the participants’ mind’s eye was an AM or a non-AM.

The square-shape of the time by time generalization matrix shown in Fig.3a showed that the task allowed the formation of stable representations associated with the figures (1 random polygon and 1 rounded shape for AMs and the same for the non-AMs) acting successfully as a localizer for the two types of memories. The bootstrapping analysis performed on a 0–2.5s time-window showed that the classifier could distinguish with a peak accuracy of 0.55 between AM and non-AM (*p* = 0.0129) in a sustained time-window (0 – ∼2.2 secs), including a late time window that is most likely related to the representation of the memory itself rather than to any perceptual features of the stimuli. In a second step, the classifier was trained during the presentation of the cue in the retrieval task and then tested on the pain decisiont task in a 1sec time-window starting from the onset of the face. Crucially, any consistency in the neural pattern observed across tasks would show the representation of the memories. The bootstrapping analysis revealed a significant cluster (*p* = 0.032) in a sustained time-window (0.6–1 secs) showing evidence for the online reactivation of the memory in preparation of the empathy judgement with a peak accuracy of 0.53. The result of the classifier across tasks is shown in Fig.3b.

### 3.3. fMRI results

Fig.4 shows masked clusters resulting from the whole-brain analysis.

The contrast AM>nonAM for the analysis time-locked to the onset of the context revealed a significant FWE corrected (p <. 05) cluster with a peak in the Precuneus, BA 7, MNI: [3 −64 38], (150 voxels), t(27) = 5.76 p = 0.001, in the Superior Parietal Lobule, BA 7, MNI: [−36 −58 59], (114 voxels), t(27) = 4.42, p = 0.003 extending to the Inferior Parietal Lobule (BA 40) and in the Superior Temporal Gyrus (BA 39). This contrast also revealed a cluster with a peak in the Posterior Cingulate, BA 23, MNI: [3 −28 26], (62 voxels), t(27) = 4.71, p = 0.038. Masked clusters showing greater activation for AM as compared to non-AM are depicted in Fig.4a. The opposite contrast did not reveal any significant FWE corrected cluster.

Fig.4b shows the result of the contrast AM>nonAM for the analysis time-locked to the onset of the face. Greater activation for AM as compared to non-AM was observed in a significant FWE corrected cluster (p < .05) in the Superior Frontal Gyrus, BA 10, MNI: [−18 62 23], (66 voxels), t(27) = 5.49 p = 0.024, and in a cluster in the Inferior Parietal Lobule, BA 39, MNI: [−36 61 41], (75 voxels), t(27) = 3.67, p = 0.014. This specific region of the IPL is part of the functional fractionation of the TPJ and is considered as part of the core network of the theory of mind (Schurz et al., 2014). The opposite contrast revealed greater activation for non-AM than AM in a significant FWE corrected cluster in the parahippocampal gyrus (PHG), BA 36, MNI: [−18 −16 −22], (169 voxels), t(27) = 6.22, p < 0.001.

## 4. Discussion

In the current study we recorded EEG and hemodynamic activity from two independent samples of participants to investigate whether AMs are reactivated in the service of empathy. The present results from two independent experiments provide behavioral, electrophysiological and fMRI evidence in support of a direct engagement of AM reactivation in empathy. We observed direct evidence for AM reactivation when participants were required explicit judgement of their empathy awareness (experiment 1) in an empathy task that activated the brain areas that are critical for both empathy and AM processes (experiment 2). Our experiments show important direct evidence on the role of AM in empathy thus providing insights into the mechanism implied by previous studies suggesting that participants’ past experiences interact with empathic abilities (Bluck et al., 2013; Gaesser and Schacter, 2014; Perry et al., 2011).

In experiment 1, EEG was recorded during a task that prompted empathy, and a memory retrieval task. Cortical EEG patterns during the retrieval task were used to probe the data from the empathy task for evidence of reactivation of AMs and non-AMs. We applied an LDA classifier, which was trained and tested across tasks, and found evidence for online memory reactivation when participants explicitly judged their empathy for others’ experiences. Participants could empathize more with people depicted in situations they had experienced themselves as compared to situations that they never experienced, as reflected in self-reported rates of participants’ empathy awareness. This behavioural result was replicated in experiment 2 showing the robustness of this behavioural evidence.

Three critical features of the study design underwrite the robustness of our findings. First, the autobiographical component of the memories used to probe empathy was unprompted in the empathy task. Participants’ AM retrieval could have no impact on the rating of their empathy awareness unless participants based their judgement on their own past experience. Therefore, the EEG evidence for reactivation of AM patterns is remarkable because participants could have relied entirely on their semantic knowledge to perform the tasks, yet the above chance performance of the classifier suggests that they did not. Second, the memory retrieval task was always performed after the empathy task to avoid that participants could be primed to specifically retrieve their own memories. Third, we used perceptually different stimuli to prompt empathy for specific contexts (sentences) and to trigger the reactivation of those episodes (shapes). This was done to avoid any overlap in the perceptual features that cued the memories in the two tasks and ensure that the classifier could only identify the neural fingerprint of the memories reactivation per se. Our results suggest that memory retrieval and empathic processes operate within the same time-frames. The EEG pattern classifier approach has been successfully adopted to differentiate between the retrieval of perceptual and semantic content of an episodic memory (Linde-Domingo et al., 2019) and in different mechanisms of memory (Jafarpour et al., 2013). The timing of the retrieval of an AM has been shown to occur between 400 and 600 ms even when it is only spontaneously recalled (Addante, 2015; Hebscher et al., 2019). The squared shape of the time by time generalization output depicted in Fig.3 shows that the representation is stable across time (King and Dehaene, 2014). Fig.3a shows that the representation of the memories starts stabilizing between 500ms and 1sec and lasts until ∼2 secs. Fig.3b shows that the representation of the memories reactivate in the time-window when the empathy judgement was prepared.

In experiment 2, we recorded fMRI in an independent sample of participants to verify that the empathy task activated brain areas commonly associated with empathy and AM. This second experiment replicated the behavioural result obtained in experiment 1: participants reported increased empathy awareness for individuals described in contexts for which they had an associated AM. Whole-brain analyses contrasting the hemodynamic response for AM and non-AM were conducted related to the onset of the context, i.e. when participants read the sentences describing either an AM or a non-AM, and of the face, conveying painful or neutral expression. The first analysis showed activation of the Precuneus (BA 7), PCC (BA 23) and left SPL (BA 7). The second analysis activated the left SFG (BA 10) and a specific region of the left IPL, part of the functional fractionation of the TPJ (BA 39). The activation of these brain areas is consistent with previous literature showing that these brain areas underlie both AM retrieval and empathic processes (Amodio & Frith, 2006; Bernhardt & Singer, 2012; Buckner & Carroll, 2007; Frith & Frith, 2003; Spreng et al., 2008; Zaki & Ochsner, 2012).

The parietal cortex is a critical hub for cognitive, cold, empathic processes, and AM retrieval. The SPL is involved in the online maintenance of relevant information (Postle et al., 2004; Xie et al., 2019) and in the retrieval of specific AM (Addis et al., 2004). A recent study by Hebscher and colleagues, demonstrated the causal involvement of the precuneus in AM retrieval (Hebscher et al., 2020). The involvement of the parietal cortex in the retrieval of AM, and in particular of the precuneus, has been suggested to be responsible of the spontaneous AM retrieval from an egocentric perspective (Freton et al., 2014) and in flexible perspective shifting during AM retrieval (St. Jacques et al., 2017). Therefore, it ultimately contributes to the vividness of the retrieval and of constructing realistic mental images (Fuentemilla et al., 2014). The precuneus is reliably engaged in the network of brain areas underlying the understanding of others’ mind, i.e. cold empathy (Molenberghs et al., 2016). The PCC is involved in the retrieval of familiar objects and places (Burianova and Grady, 2007) and together with at least the anterior division of the precuneus, in self-referential processes (Sajonz et al., 2010).

In the whole-brain analysis contrasting the AM and non-AM related to the onset of the face, we observed the activation of a specific region of the left IPL, part of the functional fractionation of the TPJ (BA 39) and of the left SFG (BA 10). A recent meta-analysis investigating the core network of theory of mind (Schurz et al., 2014) demonstrated that, together with the mPFC, the left TPJ is a core brain area of this network (Gaesser et al., 2019). Lesion studies further support this view as damage of the left TPJ can selectively reduce theory of mind abilities but not other cognitive or executive abilities (Apperly et al., 2004; Bzdok et al., 2013; Samson et al., 2004). The activation of the left SFG was in line with the source estimation of the ERP data in experiment 1 for the same contrast and time-window. ERP studies investigating empathy for physical pain have shown that an empathic reaction, reflecting the processing of a painful experience, is expressed as a positive shift of the ERP response, compared to a neutral condition with (e.g., Meconi, Doro, Lomoriello, Mastrella, & Sessa, 2018; Sessa & Meconi, 2015) or without (Sheng and Han, 2012) relation to explicit or implicit measures of empathy. In experiment 1, we observed a positive shift in the ERPs reflecting the processing of painful when compared to neutral faces within 1 second in a cluster of centro-parietal electrodes that was estimated to be generated in the IPL and the IFG. Within the same time-window, ERPs time-locked to the onset of the faces reflecting the processing of the preceding memory showed a positive shift of the ERPs for AM as compared to non-AM. The neural source of this effect was estimated to be in the SFG. According to the multiple memory system of social cognition (MMS), prejudice and stereotyping are the result of affective and semantic associations in memory (Amodio and Ratner, 2011) resulting from autobiographical experience as well as from acquired knowledge. Studies on cross-racial empathy for pain showed that empathic responses are more natural for own-race faces or more familiar faces when compared to other-race faces (Avenanti et al., 2010; Sessa et al., 2014a; Xu et al., 2009). These ERP studies therefore provided some parallel evidence that past experiences and shared cultural background can influence empathy as they contribute to reduce the psychological distance between the observer and the target of empathy (Meconi et al., 2015). Our ERP results and the source analysis regarding the face and memory effects replicate previous ERP (Sessa et al., 2014b) neuroimaging studies on the neural correlates of empathy (Bernhardt & Singer, 2012; Fan, Duncan, de Greck, & Northoff, 2011; Lamm, Decety, & Singer, 2011) and are in line with our fMRI results obtained in experiment 2. However, it is important to mention that the area obtained from the source reconstruction in experiment 1, i.e. the SFG, seems not to fully overlap with the one observed in experiment 2 from the fMRI. It is possible that the wider cluster obtained in experiment 1 could have been due to lower precision in the source reconstruction of an ERP effect compared with source localization from the fMRI data. Equally, though it is possible that the MNI coordinates in the fMRI analysis only identify the peaks of the clusters that in fact reflect the activation of the same, larger, brain area identified from the ERP data.

In the current study, we did not observe the activation of the MTL in the contrast AM>non-AM. Notably, the participants’ AMs were on average 5 years old. A recent review (Barry and Maguire, 2019) highlighted that although memories seem to become independent from hippocampal activation with remoteness in time, the hippocampus remains involved in context/memory reconstruction (Zeidman and Maguire, 2016) even though the original memory trace is with time transferred to the neocortex. We did observe the activation of the PHG in the contrast non-AM>AM. This result was surprising and we speculate that it is consistent with the mindreading hypothesis (Gaesser, 2018) that draws on those studies with healthy participants showing the involvement of the episodic simulation in performing tasks that prompt cold empathy. Consistently, patients with medial temporal lobe lesions do not show increases in empathy when prompted to use episodic simulation to construct specific episodes of others suffering (Sawczak et al., 2019).

## 5. Conclusions

The present study provides important evidence of a re-activation of AMs in the context of empathy. However, puzzling previous evidence showing little empathy impairment in patients with amnesia opens future research question on whether memory causally drives empathy judgments. This would require future work that modulates memory retrieval in a time-sensitive manner.

## Acknowledgements

The Authors thank all the participants and Ross Wilson and Nina Salman for helping with fMRI data collection.

## Abbreviations

*AM*: Autobiographical memory;
*non-AM*: non-autobiographical memory;
*LDA*: Linear Discriminant Analysis.
*IPL*: Inferior Parietal Lobule;
*PCC*: Posterior Cingulate Cortex;
*PHG*: Parahippocampal Gyrus;
*SFG*: Superior Frontal Gyrus;
*SPL*: Superior Parietal Lobule;
*TPJ*: Temporoparietal Junction.

## Author contribution

F.M. formulated the research question, collected and analysed all the data, manually drew the ROIs around the hippocampus and wrote the manuscript. F.M., S.H. and I.A.A. designed the studies. S.H. and I.A.A. supervised the analysis and substantially contributed to the writing of the manuscript. C.F. helped with the analysis of the fMRI data. B.S. supervised the fMRI analysis and the drawing of the ROIs. J.L.D. and S.M. helped with the classifier and bootstrapping analysis. All the authors gave important feedback and comments to the manuscript.

## Data sharing statement

The study was not formally pre-registered but the data from this research are available to view in the OSF repository: https://osf.io/9z2uf/?view_only=6b4f7e6d52a3411bbb6b4343aff79607

